# *k*pLogo: positional *k*-mer analysis reveals hidden specificity in biological sequences

**DOI:** 10.1101/102921

**Authors:** Xuebing Wu, David P. Bartel

## Abstract

Motifs of only 1–4 letters can play important roles when present at key locations within macromolecules. Because existing motif-discovery tools typically miss these position-specific short motifs, we developed *k*pLogo, a probability-based logo tool for integrated detection and visualization of position-specific ultra-short motifs. *k*pLogo also overcomes the limitations of conventional motif-visualization tools in handling positional interdependencies and utilizing ranked or weighted sequences increasingly available from high-throughput assays.

---

The specificity of many biological processes relies on the recognition of sequence motifs. Accordingly, sequence-motif analysis, including both discovery and visualization of motifs, has long provided fundamental insights into molecular biology. However, most existing motif-analysis tools have fundamental limitations.

Existing motif-visualization tools, such as WebLogo^1^, iceLogo^2^, and pLogo^3^, usually take a set of aligned sequences as input, calculate the weight (frequency or statistical significance) of each letter at each position, and generate logo plots in which letter heights are scaled relative to their weights. Because each position is considered separately, these tools are unable to model and visualize interdependence among multiple positions and thus cannot resolve motifs that overlap with each other. Moreover, these tools treat each input sequence equally and thus do not support weighted or ranked sequences, which are increasingly available from high-throughput studies, such as *in vitro* selection^4^ and massively parallel reporter assays^5^.

In contrast, existing motif-discovery tools exclusively model interdependences between neighboring letters, and some can handle weighted or ranked sequences^6,7^. However, unlike motif-visualization tools, which precisely model each position, motif-discovery tools typically ignore positional information and thus miss ultra-short motifs (with lengths 1–4 letters) or other information-poor motifs whose specificities are conferred by both sequence identity and relative position. Examples of such hidden specificity have recently been discovered by high-throughput analyses of interactions that were previously thought to be sequence-independent^4,8,9^.

Because of the limitations of existing tools and the strong synergy between motif discovery and visualization, we developed an integrated framework for sensitive detection and visualization of position-specific ultra-short motifs from either weighted or unweighted sequences. Our tool, called *k*pLogo (*k*-mer probability logo), extended the framework of pLogo (probability logo), which scales the height of each motif residue to show the statistical significance of its enrichment^3^, in two aspects. First, to support ranked and weighted sequences, *k*pLogo calculates test statistics and their corresponding *P* values using Mann-Whitney *U* tests and Student’s *t* tests, respectively. Second, in addition to testing the statistical significance for single letters at each position, *k*pLogo tests it for all short *k*-mers starting at each position (default *k* ≤ 4, allowing degenerate letters). To visualize the results, *k*pLogo generates a new type of logo plot called the *k*-mer logo, in which at each position the most significant *k*-mer is plotted vertically with the total height scaled to its *P* value (−log_10_ transformed) or test statistics, as appropriate.

To illustrate the ability of *k*pLogo to examine weighted or ranked sequences, which are currently not handled by widely used tools such as WebLogo and pLogo, we used *k*pLogo to summarize and visualize the results of a massively parallel reporter assay in which 26,438 variants of the cAMP-responsive enhancer were generated and assayed for activity^5^. In the original publication, the relative importance of each nucleotide at each position was visualized by either four bar plots (one for each of the four nucleotides) or a heatmap resembling that of **Fig. 1a** (top). Starting from the list of variant sequences and their activity scores, *k*pLogo performed Student’s *t* tests to evaluate for each nucleotide at each position whether variants with that nucleotide at that position had higher activity than the rest of variants, and generated a probability logo with more readily visible sequence motifs (**Fig. 1a**, bottom).

**Figure 1.**
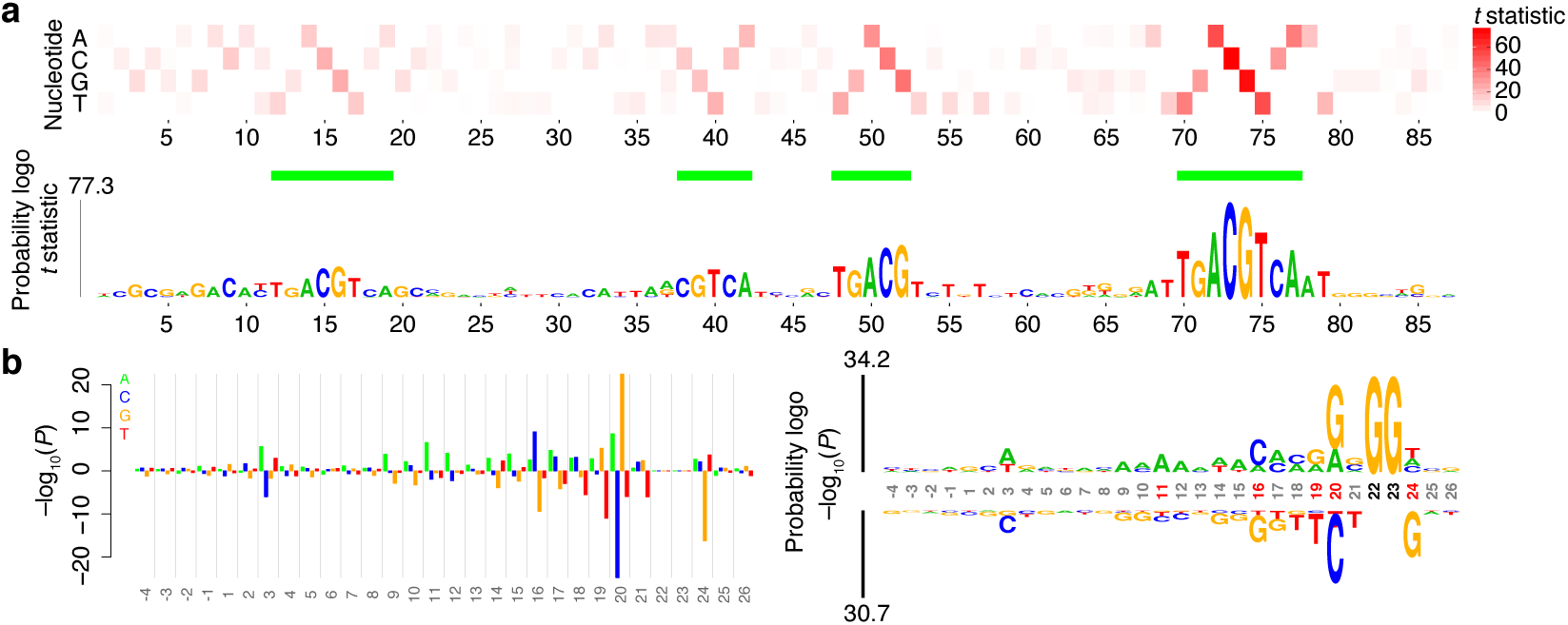
Analysis of weighted and ranked sequences. (**a**) Comparison of a heatmap (top) and a *k*pLogo probability logo plot (bottom) for summarizing results of reporter assays for 26,438 variants of the cAMP-responsive enhancer. Both formats visualize *t* statistics from Student’s *t* tests. Only the top half of the *k*pLogo probability logo plot is shown. Four consensus CREB sites are highlighted (horizontal green bars). (**b**) Comparison of a barplot (left) and a *k*pLogo probability logo plot (right) for depicting the *P* values from Mann–Whitney *U* tests of whether guide RNAs with a specific nucleotide at a specific position are more (above 0) or less (below 0) efficient than other guide RNAs. In the probability logo, positions with significant nucleotides (Bonferroni corrected *P* < 0.01) are highlighted (red coordinates), as are the fixed positions of the GG PAM (black coordinates).

We also used *k*pLogo to analyze and display the results of a study that ranked the efficiency of 1,841 Cas9 guide RNAs designed to knock out reporter genes^9^. The original publication compared the guide RNAs in the top quintile of efficiency (an arbitrary cutoff) to the rest, calculating the enrichment and depletion of each base at each position using a binomial test and then plotting the results on a bar plot resembling that of **Fig. 1b** (left). Existing tools would not have been helpful in this analysis, as illustrated by the WebLogo plot generated using the top quintile of guide RNA sequences, which did not uncover any visible signal beyond the GG protospacer-adjacent motif (PAM), which did not vary (**Supplementary Fig. 1a**). In contrast, *k*pLogo could use these ranked data without imposing an arbitrary binary cutoff by performing Mann-Whitney *U* tests to find nucleotides at each position that were associated with higher or lower efficiency. The *k*pLogo-generated probability logo was also easier to read compared to a bar plot that graphed the same results (**Fig. 1b**, left vs right). Moreover, the *k*pLogo-generated *k*-mer logo uncovered preference for not only mononucleotides but also di-, tri-, and tetra-nucleotide motifs at specific positions within the guide RNA (**Supplementary Fig. 1b**).

The utility of the *k*-mer logo for revealing ultra-short position-specific motifs was also illustrated in its analysis of the primary transcripts of human microRNAs (miRNAs). miRNAs are a class of small RNAs that direct the post-transcriptional regulation of most human mRNAs^10^. They are processed by endonucleolytic excision from stem-loop regions of primary transcripts known as pri-miRNAs (**Fig. 2a**)^8,11^. For about a decade, the only features of pri-miRNAs known to be recognized by the processing machinery were structural (e.g., pairing within the pri-miRNA stem and unstructured RNA in the loop and flanking segments). Consistent with this early understanding of the defining features of pri-miRNAs, no visible sequence signals were observed when human miRNA precursors were anchored at the sites of initial endonucleolytic cleavage and visualized by WebLogo^1^ (**Fig. 2b**), nor did motif-discovery tools such as MEME^12^ identify any significant motifs within 25 nucleotides (nt) of these cleavage sites. In reality, however, the processing machinery also recognizes position-specific ultra-short motifs. High throughput experimental analyses of many pri-miRNA variants has recently revealed four ultra-short motifs that promote pri-miRNA processing, including a UG motif at the base of the hairpin (14 nt upstream of the 5′-cleavage site), a UGU motif in the apical loop (22-nt downstream of the 5′-cleavage site), a mismatched GHG motif (3 bp downstream of the 3′-cleavage site, H = non-G), and a CNNC motif (17 or 18 nt downstream of the 3′-cleavage site, N = any nucleotide) (**Fig. 2a**)^4,8^. The reason that these motifs had not been identified earlier is that they are partially redundant with each other and most important for pri-miRNAs with suboptimal structural features^8^. Therefore, although they are enriched in human pri-miRNAs and have been preferentially conserved in evolution, most of the motifs are each found in fewer than half of the human pri-miRNAs^4,8^.

**Figure 2.**
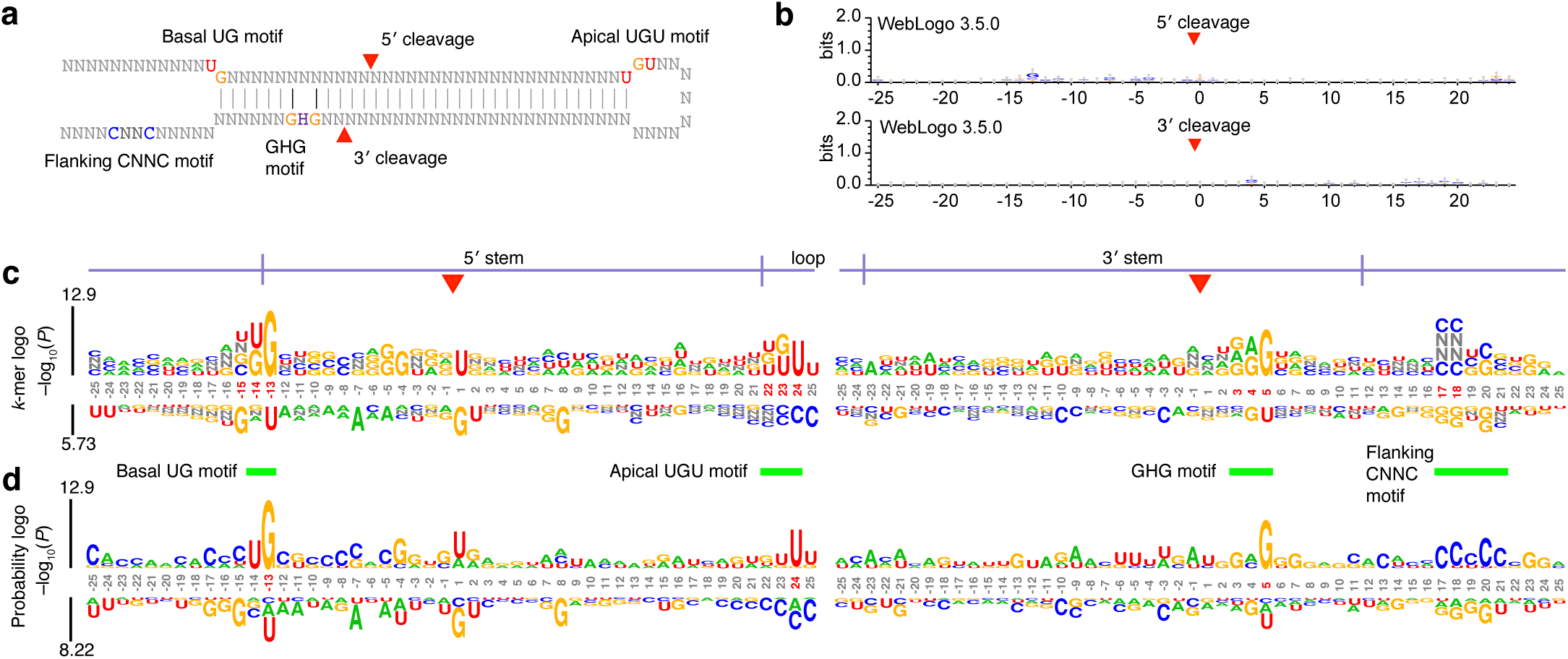
Analysis of unweighted sequences from human miRNA hairpins. (**a**) Diagram of a model miRNA hairpin indicating the two cleavage sites and four motifs. (**b**) WebLogo (version 3.5.0) output using 50 nt centered on each cleavage site. (**c**) *k*-mer logo generated by *k*pLogo using 50 nt centered on the 5′ cleavage site (left) and the 3′ cleavage site (right), with *k* ranging from 1–4, allowing the degenerate nucleotide N. The *k*-mer logo shows the most enriched (above the coordinates) and most depleted (below the coordinates) *k*-mer starting at each position, with the option of showing instead those ending at each position (**Supplementary Fig. 2**). *k*-mers read from top to bottom. Positions with significant enrichment or depletion (Bonferroni corrected *P* < 0.01) are highlighted (red coordinates). (**d**) Probability logos generated by *k*pLogo. Nucleotides are scaled by statistical significance (*P* value) and then stacked on top of each other.

In sharp contrast to existing tools, *k*pLogo identified all four of these “hidden” motifs *in silico* starting from just the known human pri-miRNA sequences, without considering any of the experimental results (**Fig. 2c**, highlighted by red coordinates and green horizontal bars). Moreover, no other motif was found to have a *P* value <0.01 (**Fig. 2c**, Bonferroni corrected, one-sided binomial test), which demonstrated high specificity. Notably, the *k*-mer logo correctly identified the CNNC motif starting at either position 17 or position 18 (**Fig. 2c**), which would have been incorrectly interpreted as a single CCNCC motif from other types of logos, such as the probability logo shown in **Fig. 2d**, thereby demonstrating the ability of *k*pLogo to discover and visualize overlapping motifs.

By default, *k*pLogo assigns each *k*-mer to the position of its starting (i.e., most 5′ or most N-terminal) letter and compares the significance of its enrichment to that of all other *k*-mers assigned to the same position. *k*pLogo also allows the option of assigning each *k*-mer to the position of its end letter. Comparing these two schemes using pri-miRNA sequences shows that known motifs sharing a start or end position with a stronger motif can be masked out in one scheme or the other but that the strongest motif among all overlapping ones was identified in both schemes (**Supplementary Fig. 2**).

Running *k*pLogo on pri-miRNA sequences from other bilaterian species, for which no high-throughput functional data were available, often uncovered motifs resembling those observed in human, as might have been expected from previous analyses examining these motifs in other species (although *k*pLogo found the motifs in an unbiased analysis, whereas the previous analyses searched specifically for the motifs)^4,8^. These previous analyses also found that none of the four motifs were present in nematodes miRNAs^4,8^, consistent with the observation that nematode pri-miRNAs are typically not processed when ectopically expressed in human cells^4^. To search for motifs that might facilitate the processing of pri-miRNAs in nematodes, we ran *k*pLogo on a set of 95 *Caenorhabditis elegans* pri-RNAs that had been previously curated to remove paralogous sequences. The two most prominent motifs were on opposite strands and together formed a paired CC/GGNG motif in the vicinity of the human mismatched GHG motif (**Supplementary Fig. 3**).

In summary, *k*pLogo streamlines sensitive motif discovery with logo-type visualization and enables the discovery of motifs missed by existing tools as well as the generation of sequence-logo plots for ranked or weighted sequences. With the increasing use of high-throughput sequencing for quantitative measurement of a large number of sequence variants, tools like *k*pLogo will have an expanding role in the discovery and interpretation of patterns hidden in biological sequences. *k*pLogo and a detailed manual for its implementation can be accessed at a simple webserver at http://kpLogo.wi.mit.edu, which generates frequency, information-content, and probability logos in addition to *k*-mer logos. Source code is freely available at https://github.com/xuebingwu/kpLogo under the GNU General Public License (GPL).

## ACKNOWLEDGEMENTS

We thank Kathy Lin for comments on the manuscript, Jeff Morgan for testing the server, and the Information Technology department at Whitehead Institute for Biomedical Research for hosting the server. X.W. is a Helen Hay Whitney Foundation Fellow. D.B. is an investigator of the Howard Hughes Medical Institute.

### AUTHOR CONTRIBUTIONS

X.W conceived of and designed the study, developed the tool and the webserver, performed the analyses, and wrote the initial draft of the manuscript. D.B. supervised the project. Both authors revised the paper.

### COMPETING FINANCIAL INTERESTS

The authors declare no competing financial interests.

---

**Supplementary Figure 1.**
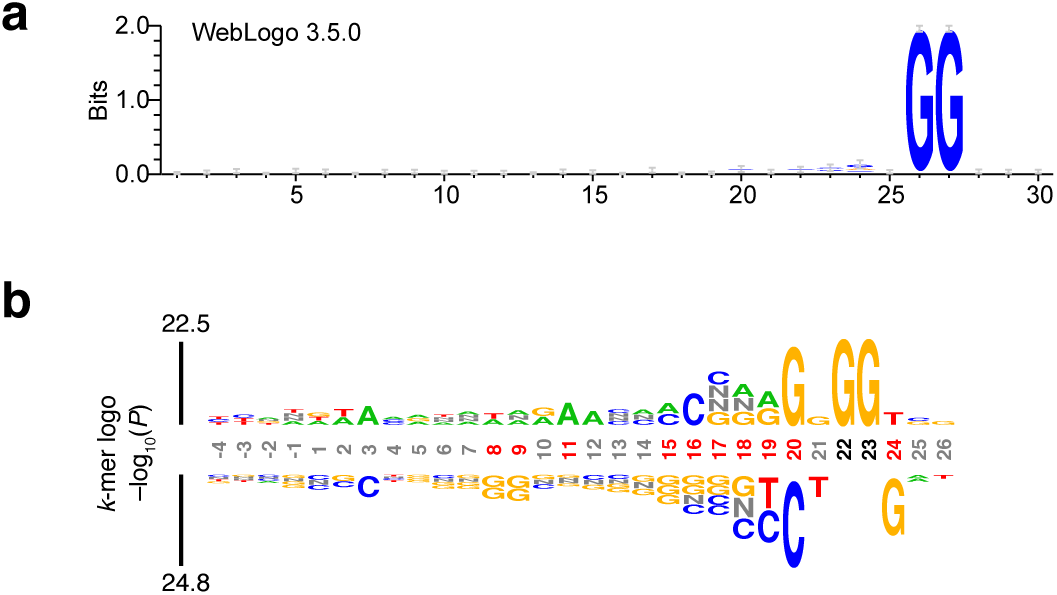
Additional representations of the ranked guide RNA results analyzed in **Fig. 1b**. (**a**) Results of running WebLogo on the guide RNAs in the top quintile of efficiency. (**b**) *k*-mer logo generated by *k*pLogo using the ranked list of 1,841 guide RNA sequences.

**Supplementary Figure 2.**
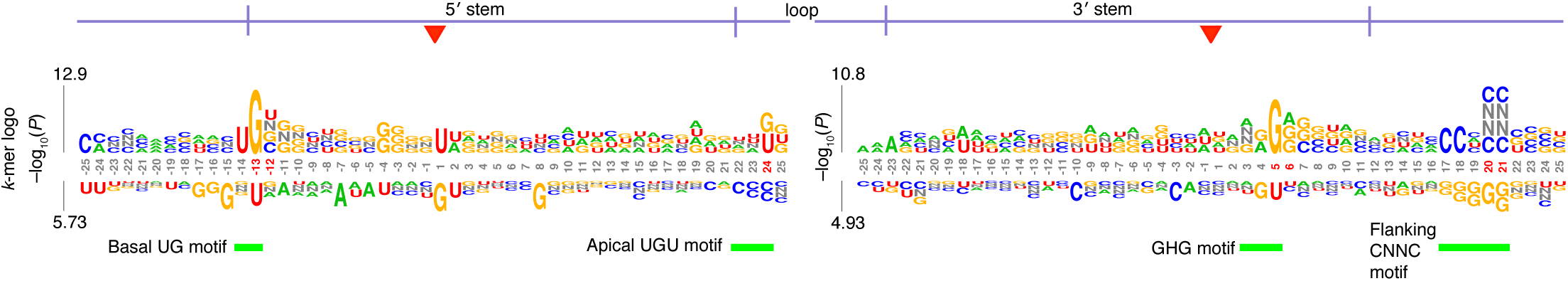
*k*-mer logo plotting the strongest *k*-mer ending at each position; otherwise as in **Fig. 2c**.

**Supplementary Figure 3.**
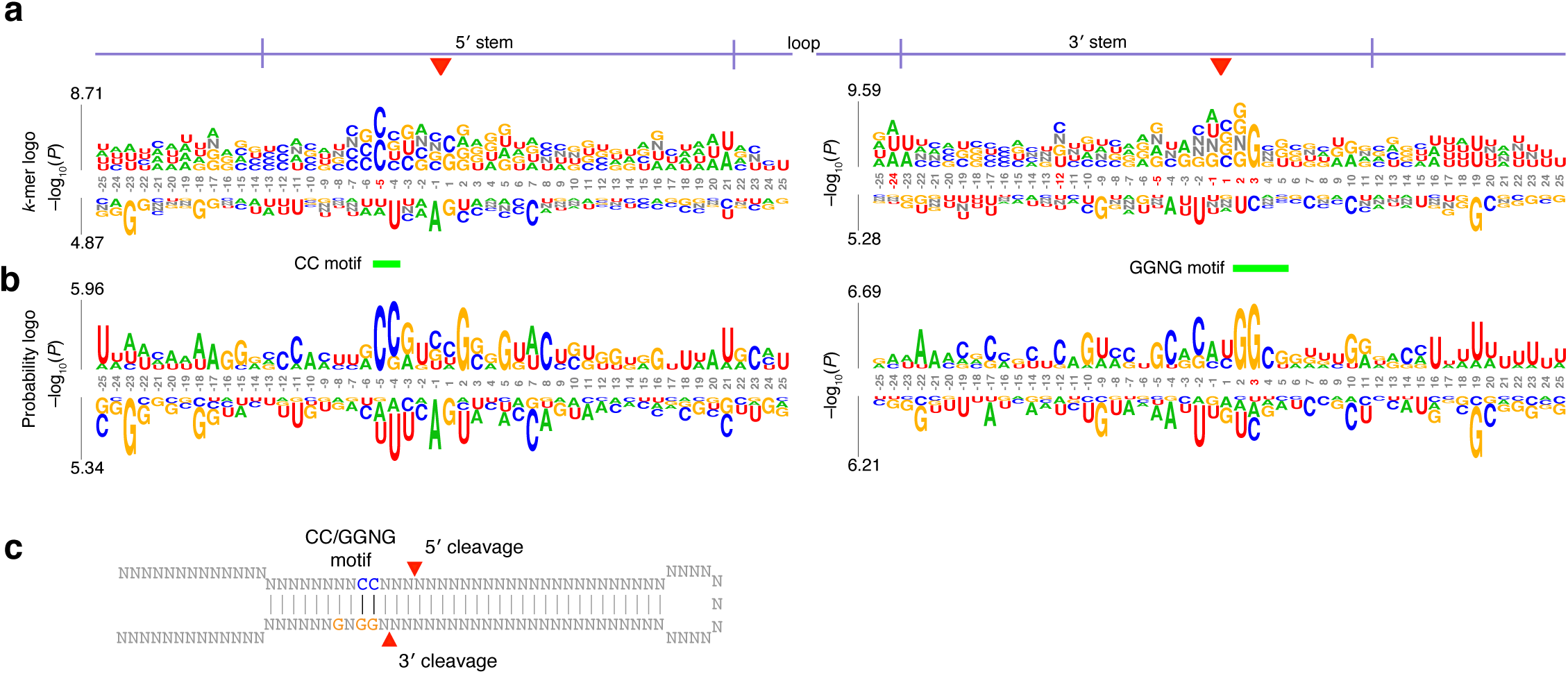
Use of *k*pLogo to identify novel motifs within miRNA hairpins of *C. elegans*. (**a**) *k*-mer logos generated by *k*pLogo using *C. elegans* miRNA hairpins; otherwise as in **Fig. 2c**. (**b**) Probability logos generated by *k*pLogo using *C. elegans* miRNA hairpins; otherwise as in **Fig. 2d**. (**c**) Diagram depicting the motifs identified by *k*pLogo using nematode miRNA hairpins.

## METHODS

### *k*pLogo tool and web server

*k*pLogo was developed in C++, and can be used via either a simple command-line interface or a web server. As input, it accepts a list of sequences of identical length, which are either unweighted, weighted, or ranked. *k*pLogo enumerates all possible *k*-mers of user-specified lengths, evaluates their presence at each position in all input sequences, and reports their enrichment and depletion at each position as determined using an appropriate statistical model (described below). *k*pLogo tests all *k*-mers ranging from 1–4 letters by default and can also be configured to test *k*-mers of other lengths. Degenerate letters can be specified using the IUPAC code. In addition to probability logo and *k*-mer logo, *k*pLogo also generates logo plots for monomer frequency and information content.

### Statistical models

#### Weighted or ranked sequences

For each position and for every possible *k*-mer of user-specified size range (default from 1–4) at that position, input sequences are divided into a positive group and a negative group, depending on whether a match to the *k*-mer can be found at the specific position in the sequence. The weights in the two groups are then compared using the one-sided two-sample Student’s *t* test, or ranks in the two groups are compared using the Mann–Whitney *U* test (using a *z*-test approximation).

#### Unweighted sequences

For each possible *k*-mer of user-specified size range (default from 1–4) at each position, the one-sided binomial test (using a *z*-test approximation) is used to evaluate whether the frequency of the *k*-mer is higher or lower than expected. The expected frequency is determined using one of three background models specified by users to be either the average frequency of the same *k*-mer across all positions (default), the frequency of the same *k*-mer at the same position but in a separate set of background sequences or shuffled input sequences that preserve sequence composition, or the frequency calculated from Markov models learned from the input sequences or background sequences.

### Logo generation

In each position of a *k*-mer logo, only the most enriched *k*-mer and the most depleted *k*-mer starting at that position are shown above and below the coordinate, respectively. Each *k*-mer reads vertically from top to bottom, with total height scaled by either its test statistic (either *t* statistic or *z* score, depending on the test) or its –log_10_-transformed *P* value, depending on the user-defined preference, although in instances in which an absolute *P* value is too small to be represented in the current computer system (*P* < 10^-324^) the analysis and scaling defaults to test statistics. In a probability logo, single letters are stacked on top of each other and each scaled by associated test statistics or *P* values. Enriched/depleted letters are stacked above/below the coordinates, respectively. In both *k*-mer logo and probability logo, coordinates of positions with *P* values smaller than a specified threshold (default 0.05) after Bonferroni correction are highlighted in red. Positions for which the frequency of a single letter exceeds a defined threshold (default 0.75) are designated fixed positions, and coordinates of these positions are highlighted in black. Only the dominant letter is shown at fixed positions, and letters at fixed positions are shown at heights 10% higher than the max height of non-fixed positions.

### Data sources

Sequences and associated weights (if any) for miRNA hairpins^4^, CRISPR guide RNAs^9^, and CRE enhancer variants^5^ were obtained from the corresponding publications and are included in the *k*pLogo source code as example inputs. The sets of miRNA hairpins from human and *C. elegans* had been curated to remove closely related paralogs^4^, which reduced the chance of identifying motifs that were enriched solely due to descent from common ancestry.

